# SEC Purified Monomeric Aβ42 Produces Reproducible and Reliable Aggregation Measurements

**DOI:** 10.64898/2026.05.12.724608

**Authors:** Jhinuk Saha, Joshua Dindinger, Ayyalusamy Ramamoorthy

**Affiliations:** Department of Chemical and Biomedical Engineering, FAMU-FSU College of Engineering, 2525 Pottsdamer St., Tallahassee, FL 32310, United States; National High Magnetic Field Laboratory, 1800 E. Paul Dirac Drive, Tallahassee, FL 32310, United States; Institute of Molecular Biophysics, 91 Chieftan Way, Florida State University, Tallahassee, Florida 32304, United States; Biophysics and Department of Chemistry, University of Michigan, Ann Arbor, MI 48109-1055, United States

**Keywords:** Amyloids, SEC, Aggregation, Monomers

## Abstract

The accumulation of amyloid-beta (Aβ) plaques is a hallmark of Alzheimer’s disease (AD), with Aβ42 representing the predominant and most aggregation-prone isoform. Reliable preparation of monomeric Aβ42 is essential for investigating the kinetics and mechanisms of its aggregation into oligomers and fibrils. This study provides a direct comparison of two monomerization protocols for recombinantly expressed Aβ42: one incorporating size-exclusion chromatography (SEC) and the other relying solely on chemical denaturation, using agents such as NaOH and NH_4_OH. Aβ42 was produced in *E. coli*, purified through urea solubilization followed by HPLC, and subjected to monomerization via the respective methods. Monomeric preparations were evaluated using Thioflavin T (ThT) fluorescence to assess aggregation kinetics, TEM to detect fibrils and preformed aggregates, and NMR spectroscopy. SEC-isolated monomers displayed sigmoidal aggregation profiles in ThT assays, featuring distinct lag, growth, and plateau phases consistent with secondary nucleation-dominated models as determined by AmyloFit analysis. Increasing the initial peptide concentration resulted in higher fibril yields, which was further supported by TEM images showing extensive fibrillization following incubation. In contrast, non-SEC preparations containing pre-existing aggregates detectable by TEM and showed attenuated NMR signals, leading to impaired aggregation behavior. NaOH-denatured samples predominantly exhibited flat ThT curves, whereas NH_4_OH-denatured samples displayed extended lag phases. NH_4_OH performance better than NaOH, likely because its gradual pH neutralization reduced peptide structural perturbation. Overall, these findings demonstrate that SEC is critical for obtaining highly pure monomeric Aβ42 and improving the reproducibility of aggregation assays, highlighting the importance of standardized monomer preparation protocols in AD research.

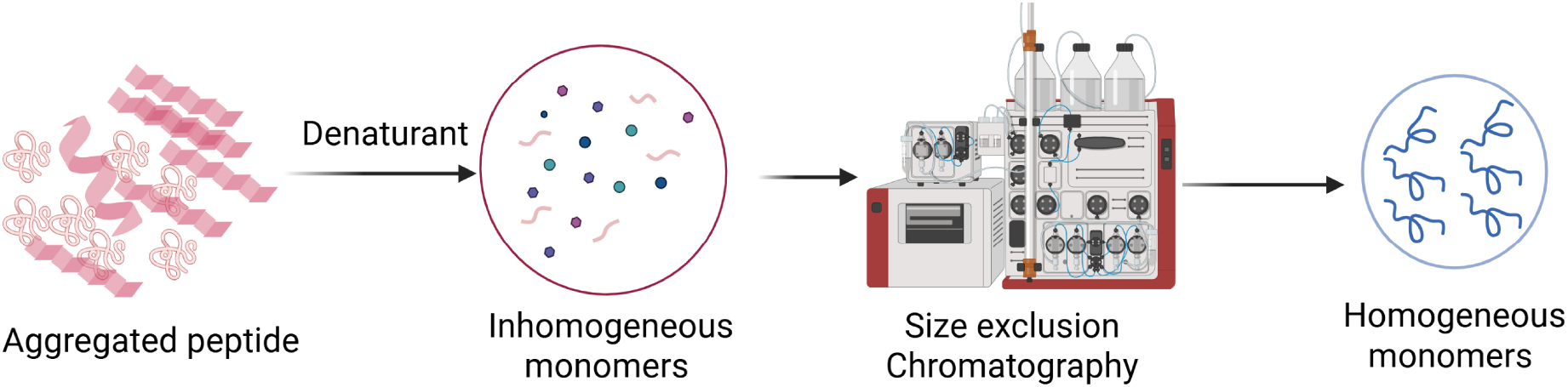

## Introduction

The accumulation of amyloid plaques in the brain, a hallmark of neurodegenerative disorders such as Alzheimer’s disease (AD), is believed to contribute to its pathogenesis. This concept underlies the amyloid hypothesis, which posits that excessive plaque buildup initiates a cascade of events leading to synaptic dysfunction, neuroinflammation, neuronal death, and ultimately, dementia.^1^ The plaques are composed of amyloid-beta (Aβ) peptides, of which Aβ42 is considered the most toxic, due to its higher propensity to aggregate.^2^ To understand the mechanism by which plaque formation takes place, much research has been devoted to studying Aβ aggregation into fibrils and oligomers.^3–5^The most common approaches for producing Aβ peptides involve synthetic or recombinant methods. The former often employs solid-phase peptide synthesis, while the latter involves expressing Aβ in a host-most commonly *E. coli*-resulting in the formation of inclusion bodies containing the peptide, which are subsequently solubilized and purified. Synthetic methods are generally more expensive; as such, many research labs prefer to purchase synthetic Aβ from suppliers rather than produce it in-house. Additionally, synthetic production of peptides labeled with isotopes such as a combination of ^15^N and ^13^C, which is required for structural studies using NMR spectroscopy, are highly expensive. Recombinant methods can also be economically burdensome, especially if purchased commercially. However, recent advances have introduced efficient and cost-effective protocols that facilitate in-house production, many of which stem from the approach developed by Walsh and co-workers.^6^ These protocols have made Aβ research more accessible and carry meaningful implications for the field.

Investigating the aggregation of Aβ requires the preparation of monomeric peptide samples. Starting with monomers is crucial for several important reasons. Physiologically, Aβ peptides are cleaved from the amyloid precursor protein (APP) by β- and γ-secretases, initially existing in monomeric state^7^. In AD and related disorders, dysregulation of Aβ production and clearance allow these monomers to accumulate and subsequently aggregate. 4 Thus, to accurately observe each step governing the aggregation process, it is necessary to begin with monomers. Furthermore, the presence of pre-existing aggregates can serve as nucleation sites, accelerating fibril formation and bypassing the rate-limiting nucleation step in kinetic studies.^8,9^ The presence of even small amounts of aggregates can also significantly impact reproducibility, complicating data interpretation and limiting the reliability of experimental results^10–13^. Furthermore, the neurotoxicity of Aβ42 is found to be fundamentally shaped by the specific biochemical environment of its initial solubilization, which dictates the subsequent neurotoxic signaling cascades. Recent research utilizing differentiated SH-SY5Y neuroblastoma cells demonstrates that solubilizing Aβ42 with either NaOH or NH_4_OH results in aggregates that are morphologically similar in size and shape, yet they trigger divergent and distinct death pathways. NaOH-prepared Aβ42 promotes a mitochondria-centric toxicity characterized by massive reactive oxygen species (ROS) production, lipid peroxidation, and calpain activation. Conversely, NH_4_OH-prepared Aβ42 initiates a GM1-independent pathway involving necroptosis-marked by the phosphorylation of RIPK3 and MLKL-alongside early-stage, caspase-independent apoptosis. While both species commonly induce endoplasmic reticulum (ER) stress.^14^. Prior work has also shown the effectiveness of NH_4_OH in monomerization of Aβ42 over HFIP treatment^15^. The aim of the present work is to compare two methods for preparing monomeric Aβ42, distinguished primarily by the use or omission of size exclusion-chromatography (SEC). Although extensive research has been conducted using both approaches, a comprehensive side-by-side comparison is currently lacking. Both techniques are used to monomerize Aβ42 produced in-house following a protocol inspired by that of Walsh et al.^6^ Details of the Aβ purification procedure are provided in the Methods section, followed by a description of the steps taken to obtain monomers with and without SEC. The resulting monomers are subjected to a series of biophysical and biochemical characterizations, including Thioflavin T (ThT) fluorescence assays, Transmission Electron Microscopy (TEM), and Nuclear Magnetic Resonance (NMR) spectroscopy.

## Experimental section

### Materials

All reagents and solutions were prepared using analytical-grade chemicals and MilliQ water unless otherwise stated. All chemicals were bought from Millipore Sigma (Burlington, MA), Fisher Scientific (Waltham, MA) or VWR International (Radnor, PA). Sodium hydroxide (NaOH), potassium hydroxide (KOH), hexafluoroisopropanol (HFIP), and ammonium hydroxide (NH_4_OH) were purchased from Millipore Sigma. Recombinant Aβ42 peptide was expressed in *E. coli BL21DE3 pLysS cells* bought from New England Biolabs (Ipswich, MA), Aβ42 plasmid was bought from Addgene (Watertown, MA) and purified in-house following a protocol adapted from Walsh et al.^6^

### Expression and Purification of Aβ42

Aβ42 was recombinantly expressed in *Escherichia coli* using cryostock cells encoding the Aβ42 construct. Bacterial cultures were grown in LB medium (tryptone, yeast extract, and NaCl) supplemented with ampicillin as the selective antibiotic. Protein expression was induced using isopropyl β-D-1-thiogalactopyranoside (IPTG). An overnight starter culture was used to inoculate larger-volume LB media, which were incubated at 37 °C with shaking until the optical density at 600 nm reached 0.6-0.8. IPTG was then added, and cultures were grown overnight. Cells were harvested by centrifugation at 4 °C and resuspended in 20 mM Tris, 1 mM EDTA. Cell lysis was carried out by sonication on ice, followed by centrifugation to isolate inclusion bodies.

Inclusion bodies were washed multiple times with 20 mM Tris, 1 mM EDTA and subsequently solubilized using 4 M urea prepared in 20 mM Tris (pH 8.0). Insoluble material was removed by centrifugation, and the clarified supernatant was filtered through a 0.2 µm PVDF membrane. Samples were aliquoted and stored at −80 °C prior to further purification. Aβ42 was further purified by high-performance liquid chromatography (HPLC) using a Thermo Scientific Vanquish™ system. Peptide-containing fractions were collected and lyophilized for downstream use. Optical density and peptide concentration measurements were performed using a Nanophotometer (NP80, IMPLEN).

### Aβ42 Monomerization via SEC

For the SEC monomerization method, samples were processed on an ÄKTA Pure chromatography system (Cytiva) equipped with a size-exclusion column (Superdex 75 Increase 10/300 GL). Denaturing agents used prior to SEC included sodium hydroxide (NaOH), potassium hydroxide (KOH), hexafluoroisopropanol (HFIP), and ammonium hydroxide (NH_4_OH), all of analytical grade. Samples subjected to SEC were pre-treated using distinct protocols aimed at achieving the highest concentration of monomers. The first method involved taking the HPLC fraction in acetonitrile and loading it into the SEC column immediately afterwards. All other treatments involved denaturing lyophilized peptides to disturb the non-covalent interactions stabilizing aggregates. The denaturing agents used in this study were sodium hydroxide (NaOH), potassium hydroxide (KOH), hexafluoro isopropanol (HFIP), and ammonium hydroxide (NH_4_OH). Details on preparing the column are provided in the Supporting Information.

The NaOH treatment involved adding 475 μL of DI water to a lyophilized peptide sample, followed by vortexing until the powder appeared fully dissolved. Next, 50 μL of 2 M NaOH was added to the solution and mixed by pipetting up and down. The tube was sealed tightly with parafilm and then incubated for ten minutes in a bath sonicator at room temperature. The sample was then transferred to a 1 mL syringe and slowly injected into the AKTA system via the sample injection valve. After placing the collection tubes onto the AKTA rotor, the SEC protocol was initiated. Over approximately 40 minutes, the sample eluted into distinct peaks corresponding to oligomers, monomers, and fragments, as shown in more detail in the Results section. The concentration of monomers eluted into the collection tubes was determined using UV-Vis spectroscopy. The KOH treatment followed the same procedural steps as the NaOH method. The NH_4_OH treatment involved adding a 10% (v/v) NH_4_OH solution in H_2_O to the lyophilized peptide sample, followed by incubation for 1.5 hours, and subsequent addition of NaPi (sodium phosphate) buffer. Subsequently, the sample was loaded onto the ÄKTA system using the same procedure as employed for the NaOH treatment. The spectra were processed in Origin and normalized to per milligram of Aβ42 for comparison analysis.

Prior to each purification, the column was thoroughly cleaned first with a 30 mL of 250 mM NaOH solution to remove any aggregates or peptide adhered to the column, followed by a 90 mL wash with MilliQ water. Immediately before sample injection, the column was equilibrated with 30 mL of sterile-filtered sodium phosphate buffer (pH 7.4).

### Aβ42 Monomerization without SEC

For NaOH treatment, 1-2 mg of lyophilized peptide was dissolved in DI water followed by addition of NaOH to a final concentration of 50 mM. After sonication, sodium phosphate buffer was added to give a final phosphate concentration of 200 mM, and the solution was adjusted to approximately physiological pH (7.4) with phosphoric acid prior to concentration determination by UV-vis spectroscopy. For NH_4_OH treatment, 1-2 mg of lyophilized peptide was dissolved in aqueous ammonium hydroxide to yield an effective ammonia concentration of ∼4% (NH_3_, w/v equivalent). The solution was subsequently titrated with phosphoric acid to pH 7.4 before UV-visible spectrophotometric analysis. An identically prepared peptide-free blank was used for background correction for both type of preparations.

### ThT Fluorescence Assays

Thioflavin T (ThT) fluorescence assays were conducted using a BioTek Synergy H1 microplate reader. Monomeric Aβ42 obtained from each preparation method was subjected to ThT fluorescence assays following concentration determination via UV-Vis spectroscopy. To minimize premature fibril formation, monomer samples were kept on ice prior to use. For the monomers prepared using both methods, ThT assays were conducted at four different concentrations: 5, 10, 15, and 20 µM in 10 mM sodium phosphate buffer at pH 7.4. For each concentration, triplicate samples were prepared in a final volume of 50 µL. Prepared samples were loaded onto a 384 well plate and incubated in the ThT plate reader for 20 to 48 hours, depending on the trial. The plate reader was set to 37 °C with no shaking. The data collected from the reader was exported into an excel spreadsheet and subsequently analyzed using ORIGIN software.

### Transmission Electron Microscopy

Transmission electron microscopy (TEM) images were obtained using a Hitachi HT7800 transmission electron microscope. Peptide samples were prepared at concentrations of 10, 15, and 20 μM in 10 mM sodium phosphate (NaPi) buffer (pH ∼7.4) and incubated at 37 °C for either ∼22 h or, for NH_4_OH-treated samples, 60 h to match the corresponding ThT assay endpoints. Following incubation, 10 μL of each sample was deposited on copper TEM grids and allowed to air-dry for approximately 15 min, after which the grids were stained with Uranyless and dried for an additional 15 min prior to imaging. Non-incubated peptide samples derived from both SEC-based and denaturant-based monomerization protocols were also imaged to assess initial aggregation states. TEM grids were systematically examined for fibrillar and oligomeric structures. Representative images are shown below in the Results and Discussion section and also in the Supporting Information.

### Circular Dichroism Spectroscopy

Circular dichroism (CD) measurements were performed on Aβ42 samples treated with 4-10% (v/v) NH_4_OH. Following NH_4_OH treatment, samples were prepared either without further purification or after size-exclusion chromatography (SEC), as indicated. For CD experiments, Aβ42 samples were adjusted to a final peptide concentration of 20 µM in 10 mM sodium phosphate buffer (pH 7.4). Far-UV CD spectra were recorded using a Chirascan circular dichroism spectrometer (Applied Photophysics) with a 1 mm path-length quartz cuvette at room temperature. Spectra were collected over the wavelength range of 190–260 nm.

### NMR Spectroscopy

All NMR experiments were carried out using a 700 MHz Bruker NMR spectrometer equipped with a cryoprobe.

## Results and Discussion

### Effect of denaturant on Aβ42 monomer recovery in size exclusion chromatography

The SEC profiles of the effects of different denaturing agents used for monomerizing the Aβ42 peptide are shown in Figure 1. For most denaturing reagents, the first elution peak within 8-10 mL retention volume was dominant, indicating a high abundance of aggregated species. The second peak, observed between approximately 12 and 15 mL for all conditions corresponds to monomeric Aβ42. Among the tested agents, NH_4_OH produced the highest monomer yield, followed by KOH and NaOH. The sample directly obtained from HPLC fractionation, which contained approximately 55% acetonitrile, was also subjected to SEC without the intermediate lyophilization step to remove the ACN:H_2_O solvent mixture. Compared with the other preparation methods, samples containing acetonitrile yielded the lowest monomer recovery, indicating the presence of larger aggregates in the HPLC elution samples (Figure 1(D&F)). Notably, the fraction collected after the void volume showed no significant absorbance, suggesting that the aggregates were larger than the column bed exclusion limit. These aggregates were therefore likely retained in the pre-column filter and unable to pass through the column. The final peak is attributed to Aβ42 peptide fragments and was especially prominent when using KOH, suggesting excessive denaturation and fragmentation. Consequently, although NaOH yielded fewer monomers than KOH, it was selected—along with NH_4_OH—as one of the denaturing agents for non-SEC monomerization because it produced lower levels of Aβ42 fragmentation while still providing relatively good monomer yields. Higher NaOH concentration (200 mM, 400 mM and 1000 mM) were also evaluated to improve monomeric Aβ42 yield; however, they did not result in significant increases in monomeric recovery. Instead, larger fragmentation and impurity peaks were observed (Figure S1).

**Figure 1.**
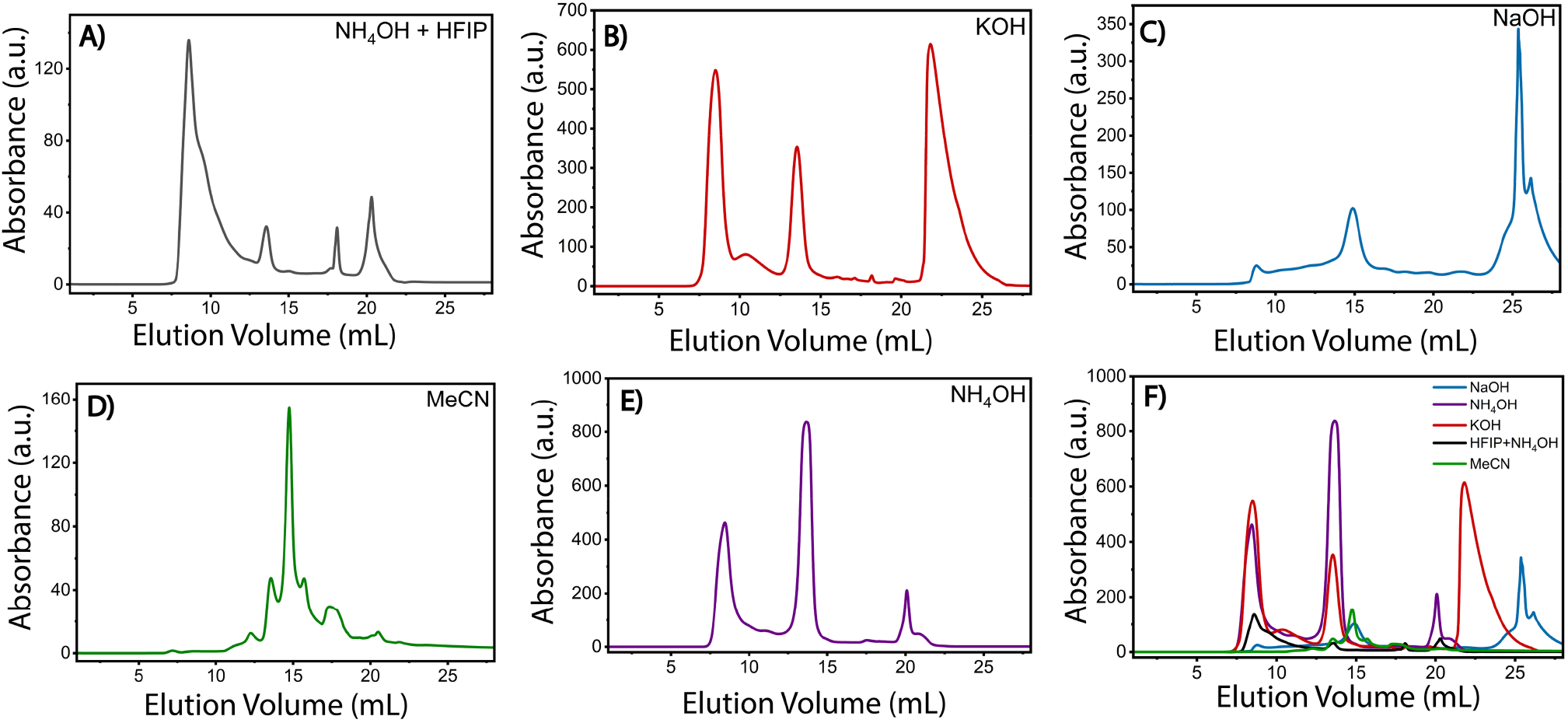
SEC elution curves showing the effects of the denaturing agents used to treat the Aβ42 peptides. (A-E) The elution profiles for HFIP+NH_4_OH, KOH, NaOH, MeCN, and NH_4_OH respectively. (F) An overlay comparison of the elution profiles for all the denaturing agents used.

### Variation in ThT Aggregation Kinetics with Monomer Preparation

The ThT experiments for each trial were conducted in the manner outlined in the methods section. Three independent ThT assays were conducted using SEC derived monomers. The average fluorescence intensity and standard deviation at each time was calculated and graphed to generate fluorescence curves for each concentration.

Aβ42 has a propensity to aggregate rapidly,^16^ and even short delays between plating and loading onto the machine can lead to the formation of some small aggregates. Nonetheless, higher starting monomer concentrations generally result in increased ThT fluorescence intensity, consistent with enhanced fibril formation as shown in Figure 2A. This observation is in agreement with prior studies demonstrating that Aβ aggregation kinetics strongly depend on the initial monomer concentration, with higher concentrations leading to more rapid aggregation.^17^ Lag times were calculated by fitting the experimental aggregation curves with a Boltzmann function, revealing a systematic shortening of the lag phase with increasing starting monomer concentration, as shown in Figure 2B. To analyze the SEC-derived aggregation kinetics, model selection was performed using AmyloFit,^18^ a software platform for the quantitative analysis of amyloid aggregation mechanisms. Following normalization of the data shown in Figure 2A, multiple kinetic models were evaluated. Although none provided a perfect fit, the two models showing the closest agreement with the experimental data are presented in Figure 2(C and D). Both involve the use of secondary nucleation dominated mechanisms^9^, with the latter being multistep. These findings are consistent with earlier reports proposing secondary nucleation dominated mechanisms for Aβ42 aggregation.^19^

**Figure 2.**
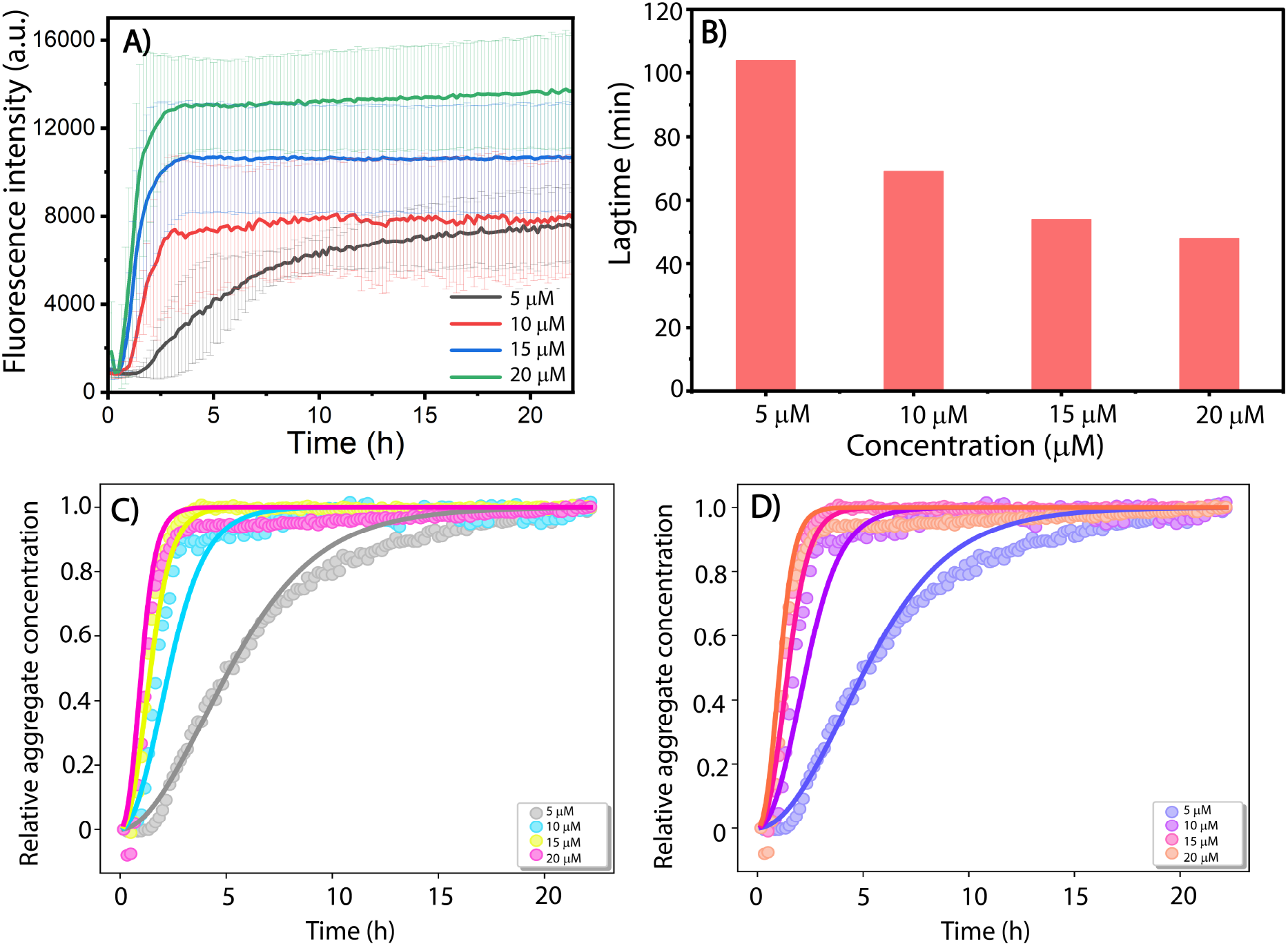
ThT assay data for the SEC derived monomers. (A) Origin graph displaying the fluorescence curves and error for the four specified concentrations. (B) Lag times calculated from the ThT curves for different concentration of Aβ42 shown in (A). (C) and (D) Using Amylofit, the secondary nucleation dominated and multistep secondary nucleation dominant fits respectively are overlayed over the fluorescence curves at different concentrations.

ThT aggregation assays were initially performed on non-SEC-derived Aβ42 monomers prepared using NaOH as the denaturing agent. This approach resulted from two observations: first, the aggregation profiles showed little to no fibril formation across most starting monomer concentrations; second, TEM analysis revealed the presence of large impurities or pre-existing aggregates in the starting “monomer” preparations. The presence of these species likely accounts for the predominantly flat aggregation curves observed in the ThT fluorescence experiments, with appreciable fibril formation detected only at 10 µM monomer concentration (Figure 3A). In contrast, Aβ42 monomers prepared using NH_4_OH as the denaturant exhibited sigmoidal aggregation behavior at 10, 15, and 20 µM concentrations as shown in Figures 3B. However, these aggregation processes occurred over a markedly extended timescale (∼20–23 h) compared with the typical 1–2 h aggregation window observed in prior assays, and the resulting kinetic profiles were not fully characteristic of canonical Aβ42 aggregation behavior.^20^

**Figure 3.**
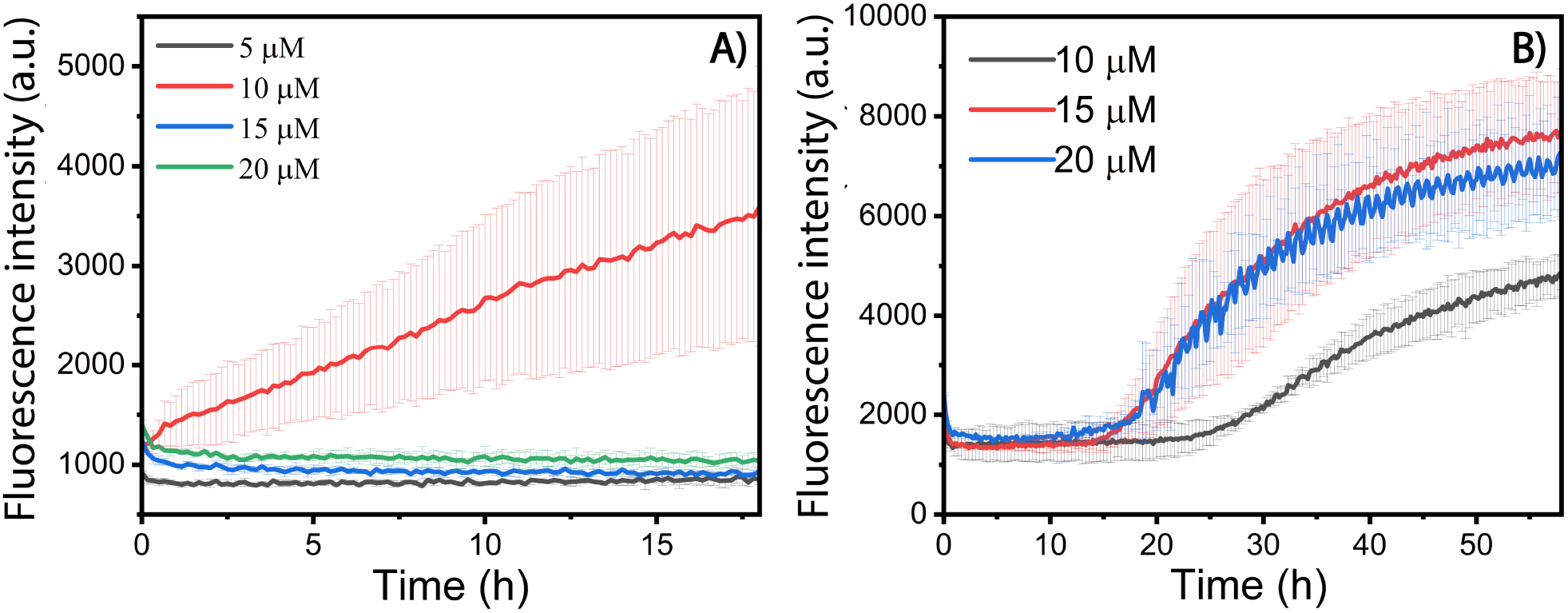
ThT assay data for the non-SEC derived monomers. Origin graph displaying the fluorescence curves and error for the specified concentrations using (A) NaOH and (B) NH_4_OH as denaturing agents.

The ThT fluorescence traces indicate that NH_4_OH may serve as a more effective denaturing agent than NaOH for aggregation studies of the Aβ42 peptide. This may be attributed to a higher initial monomer concentration in the NH_4_OH denatured, non-SEC sample. As shown by the SEC elution profiles in Figure 2, the NH_4_OH -prepared samples exhibited fewer aggregated species, whereas the NaOH-treated solution produced a significant number of aggregates. The disparity could also arise from discrepancies in pH adjustment protocols. Specifically, the NH_4_OH denatured-solution underwent a gradual change from basic to physiological pH via titration with real time pH-monitoring. In contrast, the NaOH-denatured solution underwent a more abrupt acidification. The latter protocol was based on a separate titration experiment using only buffer and NaOH, whereas attempts to perform a similar experiment with NH_4_OH resulted in precipitation, necessitating gradual adjustment instead. Although direct comparisons in the literature between rapid and gradual pH changes in Aβ42 aggregation are limited, an early study using Aβ1-28 peptides reported the formation of antiparallel β structures following a sudden pH jump from 8 to 6.5.^20^ A similar mechanism may explain the lack of further fibril formation observed in the NaOH sample as evidenced by the ThT assay reported in this study.

The fluorescence curves of the non-SEC NH_4_OH -treated samples failed to exhibit the rapid growth phase typically associated with Aβ42 aggregation. Compared to monomeric peptides isolated using SEC, the non-SEC NH_4_OH samples showed a markedly slower increase in fluorescence during the growth phase, and the time required to reach a plateau was significantly longer (Figures 2A and 3B). Furthermore, the sample prepared at 5 µM monomer concentration failed to exhibit significant ThT fluorescence, resembling the behavior of most non-SEC NaOH denatured monomers. As a result, this trace was omitted from figure 3B. The behavior of the non-SEC NH_4_OH samples is likely attributed to the presence of seeds, which may interfere with the fibrillization pathway. Overall, monomers isolated by SEC are better suited for aggregation studies, as they exhibit behavior that is more consistent with established literature and yield more reproducible results than non-SEC prepared samples.

### Aggregate Size and Population Vary with Aβ42 Monomer Preparation

Transmission electron microscopy (TEM) was used to assess fibril formation and to compare aggregate populations arising from SEC and non-SEC monomer preparation methods. TEM images of SEC-derived Aβ42 monomers (Figure 4A) showed little to no evidence of pre-existing aggregates, consistent with the observed concentration-dependent aggregation kinetics. In contrast, starting solutions of non-SEC-derived monomers prepared using NaOH or NH_4_OH (Figure 4B and 4C) exhibited a widespread presence of pre-formed aggregates and clump-like structures. The presence of these species likely contributes to the diminished extent of fibril formation observed for non-SEC “monomeric” preparations in the ThT aggregation assays shown in Figure 3.

**Figure 4.**
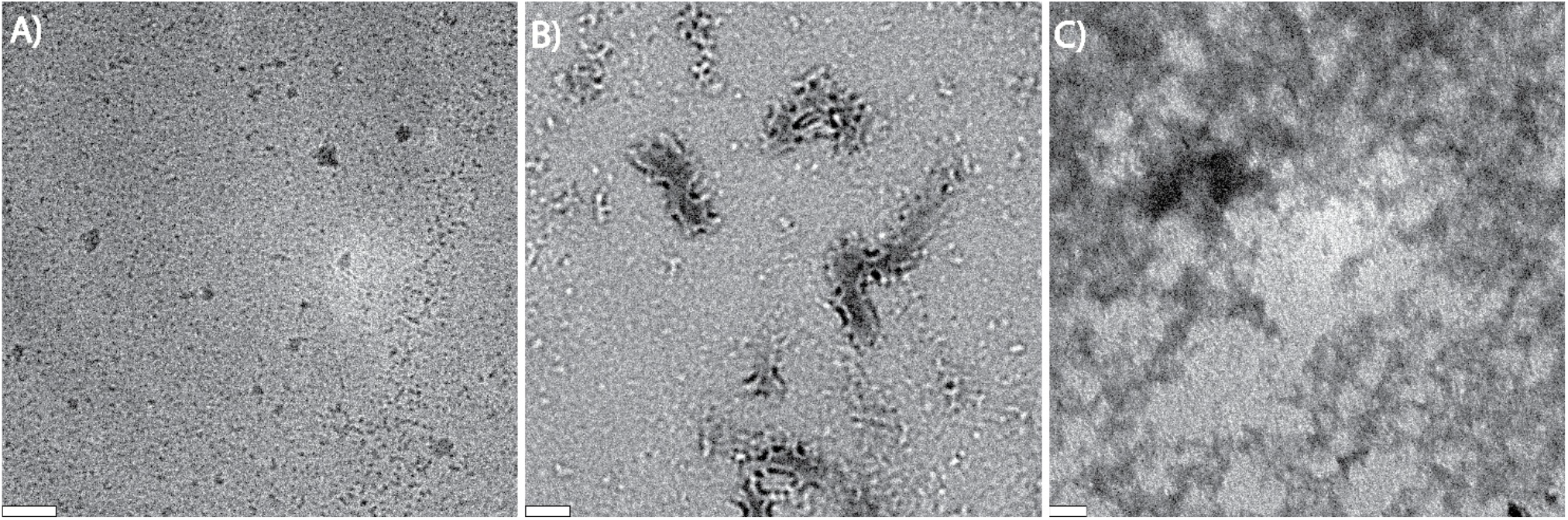
TEM images of the starting monomer solution for both monomerization techniques. (A) Monomers derived from SEC showed little to no pre-existing aggregates, as observed in this image at a 20k magnification. (B) & (C) A widespread abundance of aggregate structures observed for the non-SEC derived monomeric solutions using NaOH (B) and NH_4_OH (C) as denaturing agents. Both are at 15k magnifications. Scale bar is 200 nm.

**Figure 5.**
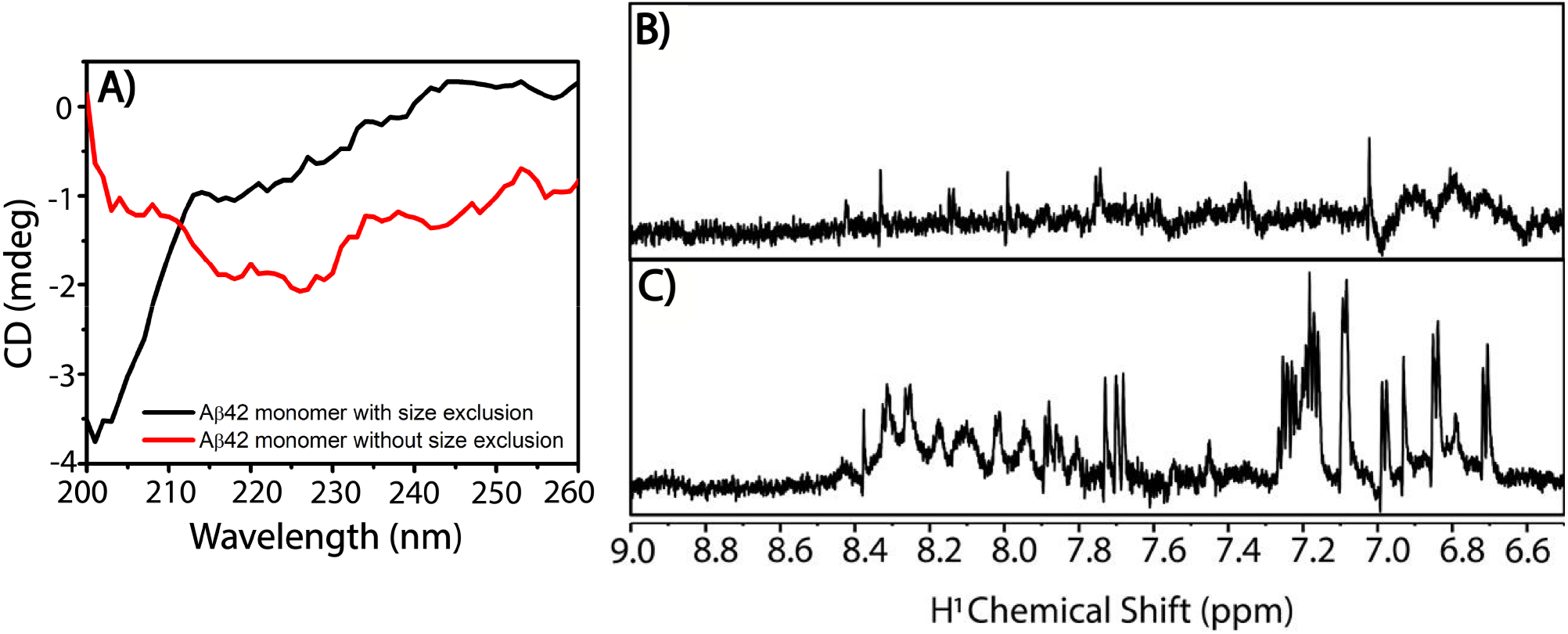
A) Circular dichroism (CD) spectra of SEC- and non-SEC–derived monomeric Aβ42 solutions in presence of NH_4_OH at 20 µM concentration in 10 mM sodium phosphate buffer pH 7.4. ^1^H NMR spectra of B) non-SEC- and C) SEC– derived monomeric Aβ42 solutions in 10 mM sodium phosphate buffer pH 7.4. Total number of scans in NMR spectrum is 256 for each spectrum and the recycle delay is 2s. Sample temperature for both CD and NMR is 298 K.

### Secondary Structure of SEC-Derived Aβ42 Monomers Differs from Non-SEC Preparations

CD spectra collected at 0 h for Aβ42 monomer preparations (20 µM) generated using NH_4_OH without SEC exhibited distinct minima at ∼230 nm and ∼218 nm, indicative of turn- and β-sheet–rich structures coexisting with monomeric Aβ42. In contrast, SEC-derived monomers showed a dominant minimum near ∼200 nm, consistent with a predominantly random-coil conformation at 0 h. These observations indicate that denaturant-based monomer preparations without SEC contain substantial amounts of heterogeneous, pre-formed aggregates that are not fully disassembled and can significantly influence subsequent aggregation kinetics.

One-dimensional ^1^H NMR spectra were collected for both SEC- and non-SEC-derived (NH_4_OH-treated) starting monomer solutions. SEC-derived samples displayed well-resolved resonances with higher overall signal intensity, consistent with a predominantly monomeric population. In contrast, non-SEC samples exhibited markedly reduced peak intensities, indicating the presence of higher-molecular-weight species that are largely invisible to solution NMR due to slow rotational diffusion and rapid transverse spin-spin relaxation (T_2_), which result in severe line broadening and signal loss. These findings are consistent with TEM observations of the corresponding starting solutions and indicate that monomerization without SEC is incomplete. The persistence of pre-existing aggregates, protofibrils, or oligomeric species likely sequesters free monomers and limits secondary nucleation, contributing to reduced fibril formation and the prolonged lag times observed in ThT aggregation assays for non-SEC preparations.

## Conclusion

This comparative evaluation establishes that size-exclusion chromatography is indispensable for the preparation of monomeric Aβ42, yielding results that are both reproducible and aligned with established aggregation behaviors. SEC effectively resolves monomers from aggregates and peptide fragments, as evidenced by elution profiles exhibiting prominent monomer peaks with NH_4_OH and KOH denaturants. Subsequent analyses supported this efficacy: TEM revealed negligible initial aggregates, NMR displayed prominent peaks indicative of monomeric species, and ThT assays produced characteristic sigmoidal curves fitting secondary nucleation mechanisms, with substantial fibril formation observed following incubation. Conversely, monomerization without SEC, dependent on denaturation alone, generated preparations contaminated with aggregates and oligomeric forms, as confirmed by extensive TEM aggregates and reduced NMR peak intensities. Such impurities likely sequestered monomers, prolonged lag phases, and impeded fibrillization, particularly in NaOH-treated samples subjected to abrupt pH shifts, resulting in aggregation patterns discordant with prior Aβ42 studies. The enhanced performance of NH_4_OH relative to NaOH in both protocols highlights the value of controlled denaturation to maintain monomeric integrity, potentially by averting antiparallel β-sheet formations associated with rapid pH transitions. These observations emphasize the inherent challenges to reproducibility in Aβ research, wherein trace aggregates can serve as seeds and confound kinetic data. Previous NMR studies have employed alkaline pretreatment methods, including NaOH treatment or dissolution at elevated pH, to denature or disaggregate Aβ and establish a more controlled initial state prior to aggregation.^21–23^ Although these approaches help reduce pre-existing aggregates and improve experimental reproducibility, alkaline treatment alone may not completely eliminate residual impurities, small preformed assemblies, or heterogeneous low-molecular-weight species that can subsequently influence aggregation pathways and structural interpretation. Advocating the use of SEC in conjunction with in-house recombinant production, as adapted from Walsh et al., would enhance both the accessibility and reliability of studies aimed at elucidating AD pathogenesis. SEC-based monomer preparation has also been employed in NMR studies to obtain a more homogeneous starting sample prior to aggregation or structural characterization. This approach helps minimize contamination from pre-existing aggregates, low-molecular-weight assemblies, salts, and other impurities that could otherwise influence aggregation behavior and complicate NMR interpretation.^24^ Study limitations encompass the single-replicate design of non-SEC ThT assays and inter-assay variability in SEC preparations, possibly arising from handling-induced premature aggregation. Future efforts could investigate gradual pH titration for NaOH in non-SEC contexts, integrate circular dichroism for structural insights, or develop hybrid approaches to optimize efficiency while preserving purity. Standardizing monomerization protocols will ultimately facilitate deeper mechanistic understanding of Aβ aggregation and inform therapeutic interventions for AD.

## Acknowledgements

This study is supported by NIH (DK132214 to A.R.). TOC figure was generated with BioRender.com. We Acknowledge Dr. Lissa Anderson (NHFML) for assisting in mass spec data collection and Dr. Benjamin Smith (NMR Facility, FSU) for assisting NMR data collection.

## Supporting Information

**Mass** spectrometry data, SEC profiles, TEM images and NMR data are included.

## Data sharing

Raw experimental data is shared at the OSF. https://osf.io/g82f9/overview?view_only=3212e2bc82bc4fafbeae9e990ca45376

