## Supporting Information for "SEC Purified Monomeric Aβ42 Produces Reproducible and Reliable Aggregation Measurements"

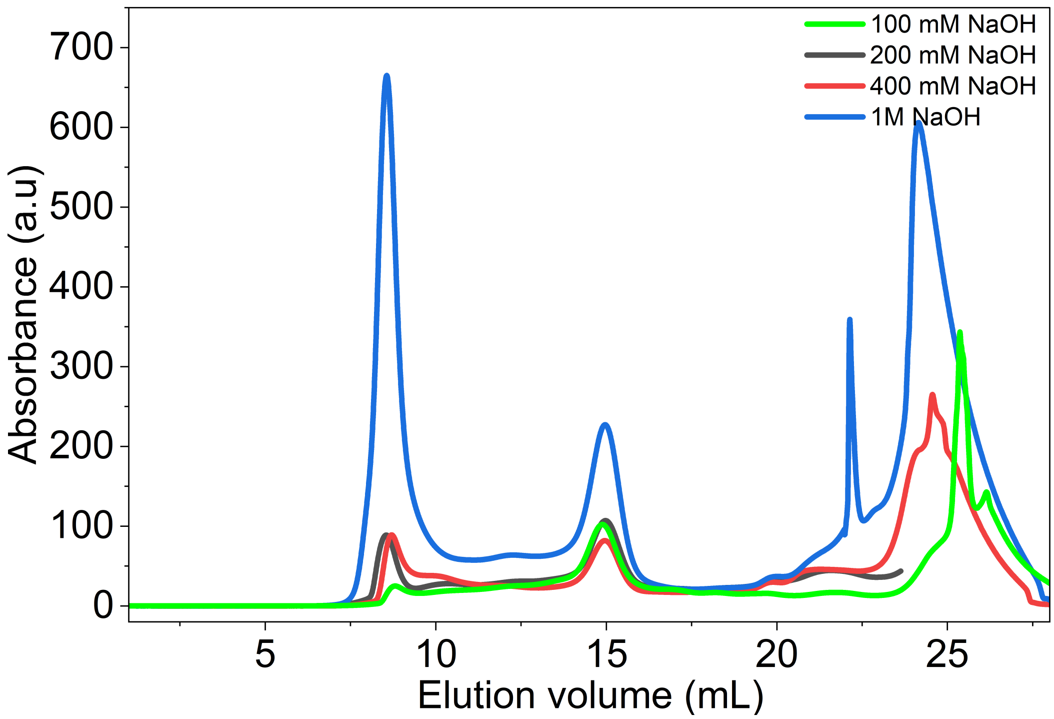


***Figure S1.*** *SEC elution curves for HPLC purified Aβ42 lyophilized and dissolved in 100 mM, 200 mM, 400 mM and 1M NaOH, and sonicated for 6-10 mins.*

**Additional TEM Images**

The SEC-derived incubated monomers, corresponding to the endpoint of previously conducted ThT assays, exhibited clear evidence of fibrillation (Figure 2) along with oligomerization and formation of other small aggregates.

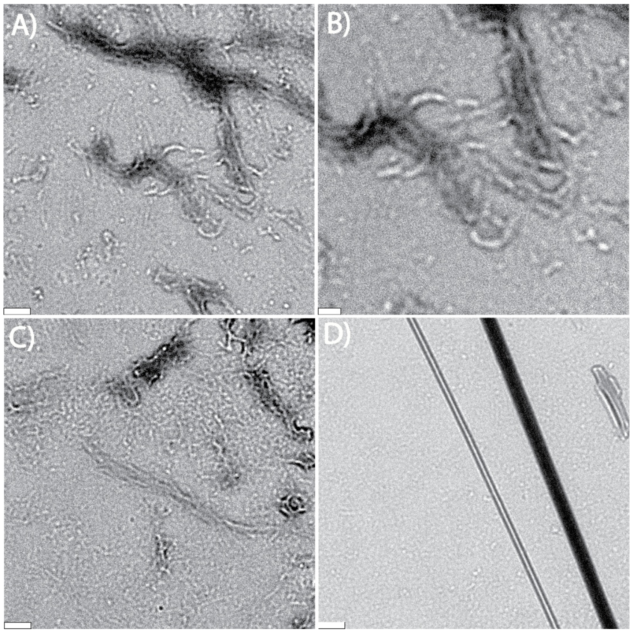


***Figure S2.*** *TEM photographs of SEC-derived Aβ42 monomers aggregating at different concentrations after a ~22-hour incubation period. (A) and (B) Fibrils found at a concentration of 10 μM at 15k and 20k magnification respectively. Scale bars of (A) and (B) are 200 nm and 100 nm, respectively. (C) Cluster of fibrils found at concentration of 15 μM at a 15k magnification. (D) Zoomed in image of singular fibrils at 15 μM and 15k magnification. Scale bar of (C) and (D) is 200 nm.*

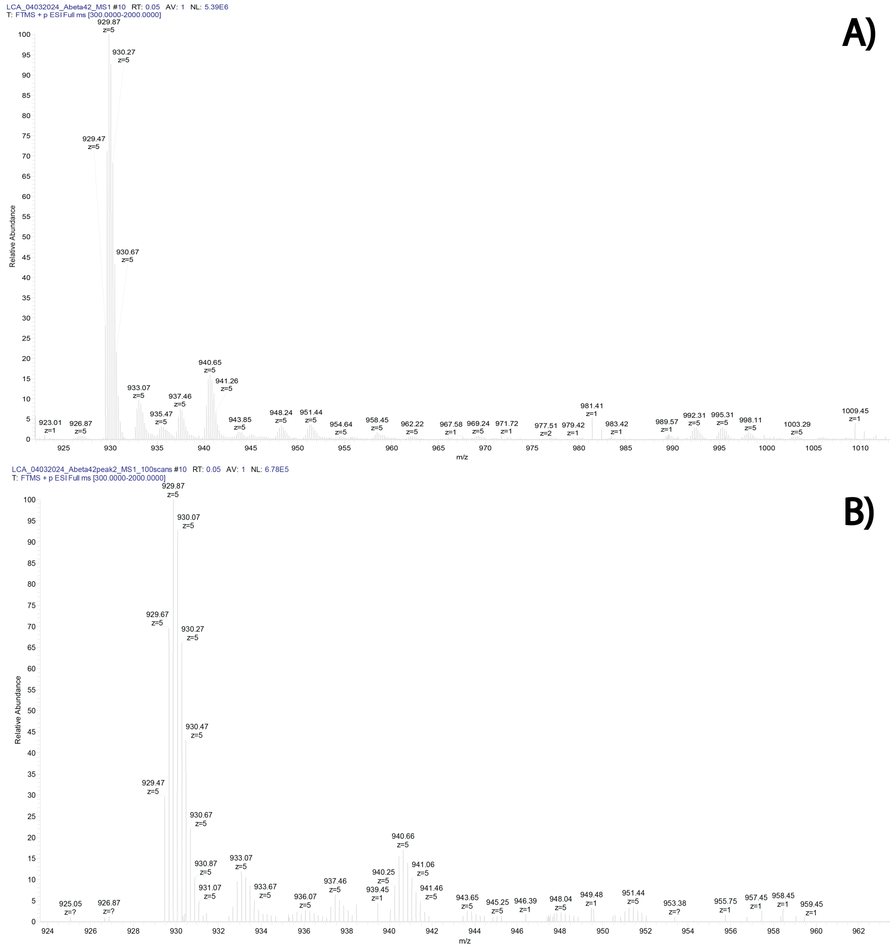


***Figure S3***. *Electrospray ionization (ESI) mass spectra of HPLC-purified recombinant Aβ42 fractions. In both spectra, the dominant signal corresponds to the +5-charge state (m/z ≈ 930), consistent with the expected molecular weight of full-length Aβ42.*

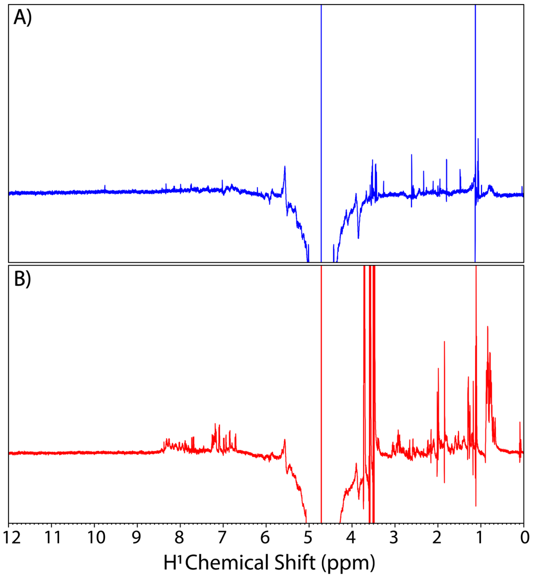


***Figure S4***. *Proton NMR spectra of Aβ monomers (a) without or (b) with SEC.*
